# Peripersonal tracking accuracy is limited by the speed and phase of locomotion

**DOI:** 10.1101/2023.04.17.537137

**Authors:** Matthew J. Davidson, Robert Tobin Keys, Brian Szekely, Paul MacNeilage, Frans Verstraten, David Alais

## Abstract

Recent evidence suggests that perceptual and cognitive functions are codetermined by rhythmic bodily states. Prior investigations have focused on the cardiac and respiratory rhythms, both of which are also known to synchronise with locomotion – arguably our most common and natural of voluntary behaviours. Unlike the cardiorespiratory rhythms, walking is entirely under voluntary control, enabling a test of how natural and voluntary rhythmic action may affect sensory function. Here, we show that the speed and phase of human locomotion constrains sensorimotor performance. We used a continuous visuo-motor tracking task in a wireless, body-tracking virtual environment, and found that the accuracy and reaction time of continuous reaching movements were decreased at slower walking speeds, and rhythmically modulated according to the phases of the step-cycle. Decreased accuracy when walking at slow speeds suggests an advantage for interlimb coordination at normal walking speeds, in contrast to previous research on dual-task walking and reach-to-grasp movements. Phasic modulations of reach precision within the step-cycle also suggest that the upper limbs are affected by the ballistic demands of motor-preparation during natural locomotion. Together these results show that the natural phases of human locomotion impose constraints on sensory function and demonstrate the value of examining dynamic and natural behaviour in contrast to the traditional and static methods of psychological science.

## Introduction

The field of active perception investigates how our experience of the world is conjointly determined by sensation and motor commands (Engel et al., 2013; Herwig, 2015; Hommel & Wiers, 2017; O’Regan, 1992; O’Regan & Noë, 2001; Press & Cook, 2015; Ridderinkhof, 2014; Schroeder et al., 2010; Witt, 2011; Witt & Riley, 2014). This approach springs from the roots of ecological psychology (Lobo et al., 2018; Witt & Riley, 2014) and in recognition of the importance of the perception-action loop, opposes the traditional separation between movement and psychophysics that dominated for many decades, and required most experiments to be performed in darkened laboratories with immobile observers. By studying perception and behaviour in more natural contexts, new insights have emerged. It has become clear, for example, that motor preparation leads to changes in sensory processing that are biassed to the current action context to enable adaptive behaviour (Bajcsy et al., 2018; Engel et al., 2013; Friston et al., 2012; Warren, 2006, 2021).

As a complement to the aims of active perception, it is also relevant to investigate the wider scope of brain-body relationships to better understand sensory processing. Most studies investigating this link have focused on the rhythmic influences of the cardio-respiratory system (Allen et al., 2022; Critchley & Harrison, 2013; Heck, McAfee, Liu, Babajani-Feremi, Rezaie, Freeman, Wheless, Papanicolaou, Ruszinkó, & Kozma, 2016; Heck, McAfee, Liu, Babajani-Feremi, Rezaie, Freeman, Wheless, Papanicolaou, Ruszinkó, Sokolov, et al., 2016; Kluger et al., 2021; Kluger & Gross, 2021; Varga & Heck, 2017) and suggest that a substantial portion of traditional task-variability in perceptual performance may be attributable to cyclic changes in these bodily signals (Azzalini et al., 2019; Klimesch, 2018; H.-D. Park et al., 2014). Here, we expand the scope of this line of work to consider another naturally occurring rhythm, focusing on how steady-state human locomotion alters sensorimotor performance.

Walking is an ideal behaviour for investigating both active perception and brain-body relationships. As with respiration, walking can be executed subconsciously, or can be voluntarily controlled (Duysens & Van de Crommert, 1998). At natural speeds, walking feels automatic, but at other speeds, it requires effortful focus. Deficits in attention or executive function can cause an abnormal gait to manifest (Yogev-Seligmann et al., 2008). Outside the clinical literature, recent work has shown that locomotion may alter sensory processing in more fundamental ways (Parker et al., 2020). Locomotion alters the excitability of sensory cortical neurons in rodents (Ayaz et al., 2013; Lee et al., 2014; Niell & Stryker, 2010; Orlowska-Feuer et al., 2022; Talluri et al., 2022), and early cortical responses in human electroencephalography (Benjamin et al., 2018; Chen et al., 2022). Intriguingly, recent research indicates that neurophysiological responses to visual information may be modulated by the presence of human locomotion (Cao & Händel, 2019; Chen et al., 2022), and behavioural responses may be modulated by the phase of the gait-cycle (Burkitt et al., 2020; Chiovetto & Giese, 2013; Matthis et al., 2017, 2018), which we explore in the current study.

We have focused on the precision of reaching movements to a moving visual target as, despite the ecological importance of interlimb coordination (Georgopoulos & Grillner, 1989; Kelso et al., 1979), relatively little is known about how steady-state locomotion might determine reach accuracy. This is surprising as many everyday activities require refined interlimb coordination while walking, such as when we reach for an item on a supermarket shelf, intercept a passing ball, or reach to shake an outstretched hand. Previous work has typically investigated reach and locomotive behaviour in isolation. When combined, a few studies have focused on grasp movements when approaching a stationary object, and analysed preferred foot-stance for postural stability (Bellinger et al., 2020; Carnahan et al., 1996; Graci, 2011; Rinaldi & Moraes, 2015; van der Wel & Rosenbaum, 2007). These tasks are valuable in identifying how balance is maintained during reach, yet by focusing on a terminating approach to a stationary object, they preclude an analysis of how rhythmic locomotion might continuously modulate sensorimotor precision in an ongoing and cyclical way. Intriguingly, a small number of studies have identified a preference for the initiation of discrete arm movements at certain phases of the step-cycle (Burkitt et al. 2020; Chiovetto and Giese 2013; Nashner and Forssberg 1986; Muzii et al. 1984). Separate investigations have identified that vestibular information is incorporated into visuomotor reach commands when reaching for a moving or displaced visual target (Keyser et al., 2017; Medendorp & Selen, 2017; Oostwoud Wijdenes et al., 2019). None of these, however, have used both free walking and reaching to the immediate peripersonal space - arguably the most ecologically relevant of dynamic reaching conditions.

For this study, we designed a task to assess peripersonal sensorimotor precision continuously during locomotion by combining a wireless virtual reality (VR) environment and the relatively new framework of continuous psychophysics (Bonnen et al., 2015, 2017; Cormack et al., 2015). While walking along a simulated path, participants made reaching movements to track continuously the 3D position of an object that changed position unpredictably. The tracking task allowed us to quantify changes in reaching precision continuously over the step-cycle as each video frame of position-tracking provides a measurement, allowing rapid collection of a dense and finely time-sampled data set. This advantage allowed us to test whether tracking responses varied over time during locomotion, and within the step-cycle, while manipulating walking speed.

We hypothesised that reach performance would improve at slower walking speeds based on results showing that a decrease in walking speeds is observed when approaching a stationary target (Rinaldi et al., 2021; Rinaldi & Moraes, 2015) and to reduce cognitive demands in dual-task walking (Al-Yahya et al., 2011; Lundin-Olsson et al., 1997; Patel et al., 2014). To foreshadow our results, we found the contrary: reach accuracy was poorest when walking slowly. We also found clear phasic modulations of both reach accuracy and reaction time over the step-cycle. Both measures changed during the swing phase of each step-cycle and in the approach to heel-strike, suggesting the continuous nature of walking imposes rhythmic demands on perception-action coupling.

## Methods & Materials

### Participants

We recruited 30 healthy human volunteers, 5 of whom were excluded for incomplete data collection resulting from hardware malfunction and signal drop-out. Our final sample of 25 volunteers were all right-handed, with normal or corrected-to-normal vision (Mean age = 22.6, *SD* = 8.9). Our sample size was selected to exceed previous samples using the continuous tracking method (typical *N* < 5, Bonnen et al., 2015, 2017) in order to compensate for the novel inclusion of our gait-based analysis. Participants received financial compensation (20 AUD per hour) or course credit, and provided written informed consent prior to the experiment. The study protocol was approved by the University of Sydney Human Research Ethics Committee (HREC 2021/048).

### Apparatus and virtual environment

Participants wore a HTC Vive Pro head-mounted display (HMD) and carried two wireless hand-held controllers. The HMD contained dual 1440 x 1600 pixel displays, each a high-resolution 3.5” (diagonal) AMOLED screen with 110 degree field of view, refreshed at 90 Hz. The HMD and hand-controllers were tracked in three-dimensional space at 100 Hz resolution, using five HTC base stations enclosing an open 4.5 x 12 metre space. The position tracking data of the HMD and dominant (right) hand were analysed, and responses were collected from the wireless controller for self-pacing the onset of each trial using a trigger button beneath the index finger. We note that independent evaluations of the HTC tracking system have placed measurement error in the millimetric range (Spitzley & Karduna, 2019), although larger errors have also been recorded, particularly when occlusion occurs between base-stations and the HMD (Niehorster et al., 2017). For our purposes, we assumed HTC tracking accuracy would be relatively constant, densely sampled the tracking station with five HTC base stations, and have focused our analyses on within participant comparisons of reach performance. Large errors in HTC tracking were rare, and when identified via visual inspection were excluded from further analysis (see ***Gait extraction from head position data***, below).

The virtual environment was designed and presented within Unity (version 2020.3.14f1) incorporating the SteamVR Plugin (ver 2.7.3; SDK 1.14.15), powered by NVIDIA Quadpro M6000 graphics (Windows 10; Dell Precision 7910). The virtual environment consisted of a large outdoor scene sparsely populated with trees, that was illuminated by simulated natural daylight. Within the environment, participants were constricted to walk within a cleared space matching the dimensions of our position-tracked physical environment. The trees, ground texture, and skybox used to create the outdoor environment were all free assets available on the Unity Asset store. To enable natural, unencumbered walking conditions, we presented the virtual environment wirelessly using the HTC Vive Wireless Adapter kit (130 gram weight).

The boundaries of the cleared walking space were identified by three-dimensional objects with the appearance of flag poles (**Figure 1a**), with the starting position of each trial indicated by a large red “X” on the ground. Trial instructions were displayed at head height at the start of each trial, adjacent to the central walking guide and target (**Figure 1b**). The walking guide was a small cube (0.1 m on a side), with a directional arrow on the superior face displaying the required direction of motion on walking trials, or a stop signal on stationary trials. The target was a sphere (radius 0.05 m), which started each trial aligned at the same depth as the walking guide. To ensure the target appeared within a comfortable reach distance, target height was calibrated to 80% of participant’s standing HMD elevation (approximately chest height) at the start of each trial.

**Figure 1.**
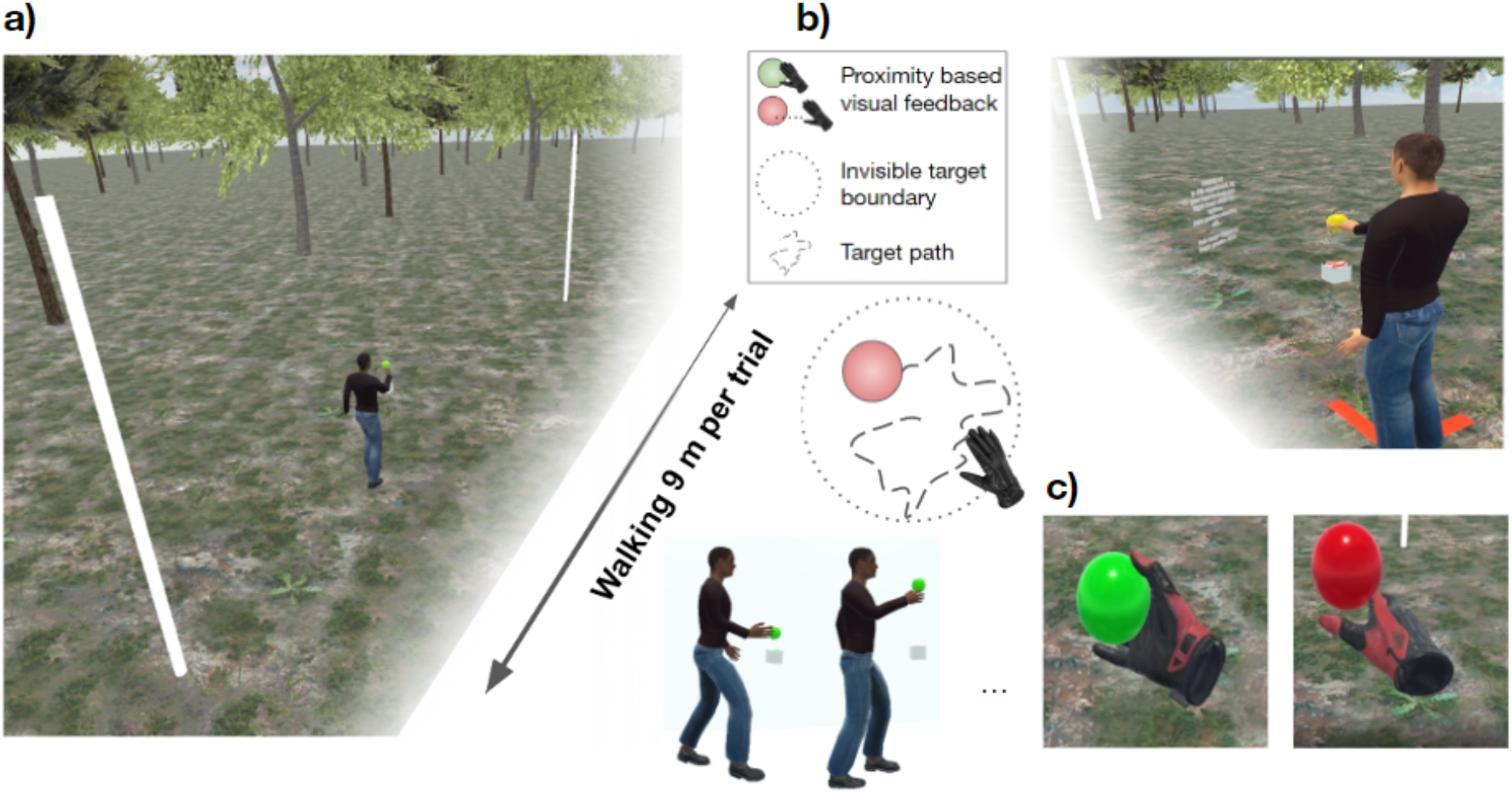
Virtual environment and first-person view of the dynamic tracking task. **A)** In each trial, participants walked along a smooth, level path in a virtual environment, **B)** while minimising the distance between their dominant right-hand and a floating sphere undergoing a random 3D walk. **C)** Visual feedback was provided in the form of a colour change (left panel) when reaching accuracy was within a predefined limit. For illustration purposes this figure shows computer-generated avatars however no avatars were used in the experiment. An example trial can be experienced from first-person view in <u>Supplementary Video 1</u>.

### Procedure and Task

Upon arrival, participants reviewed the participant information sheet, familiarised themselves with the testing environment and were given the opportunity to ask questions before providing informed consent. A brief introduction to the wireless VR apparatus, hand-controllers and battery pack was provided, before launching a practice block to expose participants to each trial combination in our design.

Our experiment employed a 2 x 2 factorial design, combining two walking speeds (Slow, Normal) with two target speeds (Slow, Fast). On all walking trials, the walking speed was set by the walking guide, which participants had to keep pace with in order to perform the task. The same 9.5 metre distance was traversed at a constant velocity, with the forthcoming walking pace indicated before the start of each trial. We set slow and normal walking trial durations at 9 and 15 seconds, resulting in walking speeds of 0.63 m/s and 1.1 m/s, respectively. These speeds were set as previous research has identified that human walkers in unconstrained, flat environments prefer an average walking speed of approximately 1.4 m/s over long distances (Hausdorff et al., 1996) and in natural environments (Hausdorff et al., 1996; Matthis et al., 2018). The slightly slower normal walking speed of 1.1 m/s was set to account for the uncertainty we expected many participants to feel when walking freely in a virtual environment.

Each experiment contained 1 practice and 8 experimental blocks, with 20 trials per block (a trial was defined as walking one length of the 9.5 m path). Each block of 20 trials contained one of the four conditions, with the order of blocks randomised per participant with the exception of the practice blocks.

The first block of each experiment was a designated practice block, during which participants experienced 2 stationary trials, followed by 2 trials of each condition in a fixed order: 1) normal Walk slow Target (nWsT); normal Walk fast Target (nWfT); slow Walk slow Target (sWsT); slow Walk fast Target (sWfT). For the remaining 10 trials, participants completed nWsT trials. For the remaining 8 blocks, the starting position was indicated by the location of the red X on the floor of the virtual environment. Participants manoeuvred themselves behind the red X to align themselves with the walking guide and were instructed to complete the task as they had practised. The target condition was displayed in text before each trial (e.g.: “On the next trial the target is SLOW, walk speed is SLOW”), and the colour of the target before trial onset was changed as a visual indicator of walk speed (slow = blue, normal = yellow).

Each walking trial began by clicking the hand-held trigger, which began the walking guide’s smooth linear motion at a constant velocity. During trials, the position of the target moved two-dimensionally in a pseudo-random manner in the fronto-parallel plane with the constraint that target position was restricted to a circle of 20 cm radius centred at the target’s starting location. The update frequency of the target movement was between 0.2 to 0.45 seconds (sampled at random from a uniform distribution with 1 ms resolution). Maximum target velocity on slow target trials was set to 0.53 m/s (distance 0 to 10.6 cm), and 0.85 m/s (distance 0 to 16.97 cm) on fast target trials. These parameters were chosen after careful pilot experimentation to induce continuous, achievable, and yet challenging motion tracking. Participants were instructed to track the target by maintaining their hand position as close to the centre of the moving target as possible, while keeping pace with the walking guide. Visual feedback was provided through colour changes to the target. The target was green if hand position was within 6 cm of the target, and red otherwise. A video containing an example trial sequence from the first-person view is located at https://osf.io/49wdt.

### Data analysis

Each trial resulted in time-series data for head, target and hand positions. We extracted step-cycle phase based on head position, and quantified task performance using the target and hand positions. Most analyses were performed in MATLAB (version 2022a) using custom scripts, and repeated-measures ANOVAs were performed in JASP 0.16.3.0. All raw data and analysis codes are available on the repository https://osf.io/jdpwc/.

### Gait extraction from head-position data

Walking results in a highly regular oscillatory pattern of movement (Hausdorff et al., 1996), with characteristic near sinusoidal changes in head position, and the vertical centre of mass over time (Gard et al., 2004; Hirasaki et al., 1999; MacNeilage, 2020; Moore et al., 2001; Pozzo et al., 1990). On the vertical axis, peaks and troughs in head position occur at the frequency of the step-cycle, with troughs corresponding to the loading phase before mid-stance – when both feet are placed on the ground (Gard et al., 2004; Hirasaki et al., 1999; Moore et al., 2001; Pozzo et al., 1990). We implemented a peak detection algorithm to single-trial time-series of vertical head position, and epoched steps based on each trough in the time-series. We excluded from all analyses any data occurring during the first two and last two steps of each trial, to allow for changes in acceleration at the endpoints of our walk trajectories. Gait percentages are typically measured from heel-strike to heel-strike of the opposite foot (step-cycle), or heel-strike to heel-strike of the same foot (stride-cycle). In our case, as we have epoched based on the loading phase of each step, heel-strike occurs slightly before the trough in the envelope of the vertical centre-of-mass (Gard et al., 2004; Hirasaki et al., 1999). In the present work, we have labelled step-cycle completion from 1-100% based on these loading-phases in our epoched time-series. Approximate locations of heel-strike and toe-off are also indicated based on prior studies which have measured foot pressure and head acceleration simultaneously (Mulavara & Bloomberg, 2002), at 10% step-cycle timing before and after mid-stance, respectively (MacNeilage & Glasauer, 2017).

We additionally performed a series of pre-processing steps which visually identified individual trials for exclusion. Each individual trial was visually inspected for data discontinuities (a result of wireless transmission delay or slip), transient spikes in error (a result of disengaging from the task), and poor gait-extraction (a result of vigorous head movement). Over all participants, an average of 7.1 trials (*SD =* 8.78) were rejected in this manner.

For our step-cycle based analyses, we additionally identified foot placement based on the direction of lateral sway (e.g., left foot stance and right foot swinging when head position is tilted to the left of centre). We then resampled the raw time-series for head, hand and target position to 100 data points (1-100% step-cycle completion). This resampling procedure is common in gait and posture research to align step and stride-cycle epochs when walking at a steady speed (MacNeilage & Glasauer, 2017), and enabled us to assess performance relative to position in the step-cycle. We note that our main results hold for the raw time-series data without resampling, owing to the highly regular step length and step duration entrained during steady-state locomotion. **Figure 4A-C** shows key points in this workflow.

### Error based on hand-target Euclidean distance

As the actual target position was known on every frame, we calculated error based on the Euclidean distance between actual target position and current hand position per time-point. For our condition comparisons, we quantified overall performance using the Root Mean Square Error (RMSE) of these distances. RMSE is common in regression analyses, and measures the average distance between the predicted values and the actual observations from the line of best fit. In our case, the observations about an idealised line of best fit correspond to the hand position and target position, respectively, and RMSE enables a quantification of error between these time-series. For our condition comparisons (**Figure 2**), RMSE was first calculated per trial, and then averaged within conditions, per participant, before performing within-participant comparisons at the group level.

**Figure 2.**
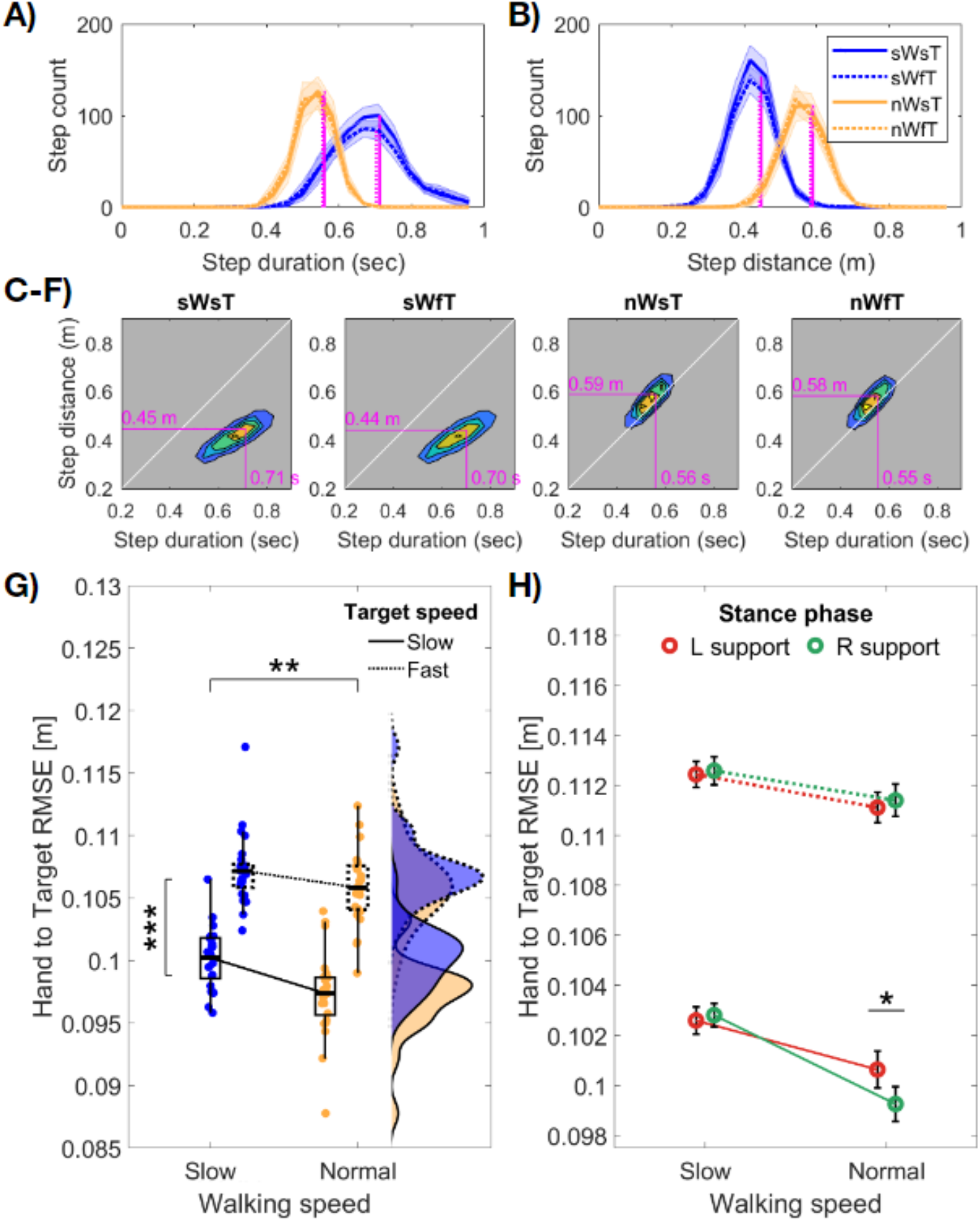
Gait parameters and average error (RMSE) per condition. In all panels, formatting for target speed (slow = solid line; fast = broken line) and walking speed (slow = blue, normal = yellow) is consistent. **A,B)** Group mean distributions of step duration and step distance per condition. Coloured lines and shading display the mean ± 1 SEM adjusted for within participant comparisons. Vertical magenta lines display the distribution means (slow targets = solid line; fast = broken line). **C-F)** Density maps of the relationship between step duration and step distance per condition. Horizontal and vertical magenta lines indicate the distribution means from a,b. Instructions to walk slowly decreased step length, and increased step duration. **G)** Average hand-target RMSE per condition, with data points displaying individual participant means. The box plots show the interquartile range and median of each condition. ** = p< .01; *** = p< .001. **H)** The data from g) when split by left or right foot support. Red colour indicates left support (right foot swinging), green colour indicates right foot support (left foot swinging). Error bars display ± 1 SEM adjusted for within participant comparisons. Reach error significantly increased when swinging the foot ipsilateral to the tracking hand, but only when tracking slow targets at normal walking speed (p < .05, FDR corrected).

### Error relative to target location

To quantify spatial changes in error, we analysed the distribution of recorded target locations on the frontal dimension (parallel to the participant’s coronal plane), as well as average RMSE per position. This analysis first tallied the distance between hand and target position at each location, before calculating RMSE based on the number of frames (i.e., duration) that the target spent at each location. For this analysis, we resampled the target-location space into a 61 x 61 grid (1 cm resolution), centred at the starting target location of each trial. At each location we quantified RMSE per trial, and then averaged within trial types, and across participants. For visualisation and analysis purposes, group-level effects were restricted to locations which contained data from all participants.

To compare the magnitude of error across target locations, we performed a non-parametric shuffling analysis to create a null distribution that removed the consistency of target location across participants. This was based on similar analyses used to compare the spatial/topographic distribution of MEG/EEG activity (Maris & Oostenveld, 2007). This analysis tests whether the observed group-average error at a specific location is greater than can be expected by chance. On 1000 permutations per participant, we changed the row and column index of each observed RMSE location, selecting a new row-column combination from two uniform distributions (with replacement). These distributions contained all possible row and column locations per participant, removing the correlation between target–location and error magnitude on each permutation. Across participants, we then compared our observed data to the 95% Confidence Interval (CI) of average RMSE from shuffled locations, and determined whether observed error was greater than expected by chance when falling beyond the bounds of this null distribution. The results of this analysis, and bounds of the null distribution are shown in Figure 3.

### Error relative to the phase of locomotion

For our step-cycle based analysis, we first quantified RMSE at all time-points within a trial, at a fixed position in the step-cycle (from 1-100%). This analysis produced an average RMSE per trial over the step-cycle, and enabled us to compare relative RMSE at different phases of locomotion, both within and across conditions.

Of central interest was whether, and when, differences in continuous reach-error would emerge over the step-cycle when comparing between target speeds and walking speed conditions. To statistically compare step-cycle based RMSE between walking speeds, we performed a series of paired-samples *t*-tests at each location in the step-cycle. We visualise significant differences between fast and slow walking speeds by highlighting step-cycle positions with *p* < .05 after false discovery rate (FDR) corrections (Benjamini & Yekutieli, 2001; Yekutieli & Benjamini, 2001).

### Cross-correlogram (CCG) and windowed Cross-correlogram (wCCG)

As a complement to distance based error, we also computed cross-correlograms (CCGs) as a proxy for reaction-time, based on the correlation between hand and target time-series. CCG functions plot the correlation between the time-series of a target and response as a function of the temporal-lag between them (e.g., Bonnen et al., 2015, 2017; Mulligan et al., 2013). To compute the CCG, we converted the position data to a velocity time-series for the hand and target on both axes (vertical and horizontal). We computed single-trial CCG functions after omitting the first and last steps (as described above), before averaging within conditions and across participants. We retained the time-lag in the peak of each CCG function as our proxy for reaction-time per condition.

In addition to single-trial CCGs, we computed short-interval windowed CCGs (wCCG) to assess reaction-time relative to position in the step-cycle. The wCCG method analyses the cross-correlation function over a short sliding window, and was originally developed to account for dependencies between behavioural time-series which may not be stable over time (Boker et al., 2002; Roume et al., 2018). This analysis followed a three step process. We computed wCCG with a short sliding window using the corrgram function (Norbert, 2007) (10 samples, approximately 110 ms duration, 1 sample overlap), and retained the wCCG function for each time-step in each trial (omitting early and late steps as described above). As a second step, each wCCG was matched to a simultaneous step within each trial, and based on the wCCG time-step was allocated to a bin representing location within the simultaneous step-cycle. For analysis, we allocated wCCG functions into quintiles, when falling within the 1-19%, 20-39%, 40-59% 60-79%, and 80-100% quantiles of a single step. As a final step, we averaged the wCCG functions within each step quintile, and compared their relative lag in peak correlation amplitude as a proxy for changes in reaction-time over the step-cycle.

### Data Visualisation

We have implemented the raincloud toolbox (Allen et al., 2019), ColorBrewer (Harrower & Brewer, 2003) and Perceptually Uniform Colormaps (https://bids.github.io/colormap/) to aid in data visualisation. The 3D avatar was placed in the Unity environment for illustration purposes only and is available from www.passervr.com.

## Results

We developed a continuous motion tracking task that varied walking speed (slow/normal) and target speed (slow/fast) in a 2×2 factorial design. Participants were instructed to minimise the distance between their dominant right hand and a moving target, while steadily walking within an enclosed 9.5 m wireless virtual reality environment (**Figure 1**). The moving target followed a pseudo-random path within a comfortable reaching distance, allowing us to assess reach accuracy in peripersonal space, and changes in task performance over the step-cycle.

### Decreased reach error at normal walking speed

As expected, our instruction to walk at a slower pace resulted in participants adopting a conservative gait, with changes evident in both step duration and step distance (**Figure 2**). A series of 2 (walking speed) x 2 (target speed) repeated-measures ANOVAs revealed that when walking slowly, step duration significantly increased (*F*(1,24) = 248.04, *p* < .001, η_p_^2^ = 0.91), while step distance decreased (*F*(1,24) = 1018.10, *p* < .001, η_p_^2^ = 0.98). On both measures, there was a significant effect of target speed (*distance*, *F*(1,24) = 17.37, *p* < .001, η_p_^2^ = 0.42; *duration*, *F*(1,24) = 14.61, *p* < .001, η_p_^2^ = 0.38) but no interaction (*p*s>.4). Post-hoc tests examining the effect of target speed indicated that the fast target speed led to decreases in both step distance and duration and thus a more conservative gait.

Having established large differences in gait parameters while walking and tracking targets at different speeds, we next quantified reach error based on the Euclidean distance between the time-series of hand and target positions (see Methods). As prior research has shown that walking speed often slows to enable prehension movements (i.e., reach-to-grasp), and that walking slowly can offset cognitive demands in dual-task scenarios, we hypothesised that our continuous reach-tracking task would be performed better at slower walking speeds. Error results shown in **Figure 2G**, however, indicate that overall, and in contrast to our hypotheses, reach error increased when walking slowly. As expected, when tracking the fast target, reach error also increased. A repeated-measures ANOVA revealed significant main effects of walking speed (*F*(1,24) = 8.66, *p* = .007, η_p_^2^ = 0.27) and target speed (*F*(1,24) = 226.26, *p* < .001, η_p_^2^ = 0.90) on reach error, with no interaction (*p* > 0.2).

We next performed an additional analysis to determine whether reach error was influenced by left or right foot support (**Figure 2H**), as prior research has indicated a task-dependent preference for ipsilateral or contralateral foot placement when reaching for a stationary object. When extending our repeated measures ANOVA to include support foot (2 x 2 x 2; walk speed x target speed x support foot) a significant three-way interaction was found (*F*(1,24) = 4.70, *p* = .04, η_p_^2^ = 0.17). Post-hoc comparisons revealed this interaction was driven by a difference in reach error between left and right foot support when walking normally and tracking slow targets (*t*(24) = 3.28, *p* = .019, adjusting for comparing a family of 28). Error was higher when standing on the left foot, and swinging the ipsilateral right foot while reaching with the dominant right hand. No other comparison based on foot support was significant.

### Reach error increases in the contralateral peripersonal space

We next investigated reach error relative to the target’s location in peripersonal space. Analysing across all trials, we observed that targets were more likely to occur centrally than peripherally (**Figure 3A**). This central tendency was expected because the random-walk algorithm reset after every trial so that all targets began at the centre of the motion boundary. **Figure 3A** also illustrates a clear symmetry of target locations around the origin which contrasts markedly with average target error per location (**Figure 3D**), which increased when reaching laterally across the body into the contralateral peripersonal space. Reaching equivalent distances in the vertical dimension did not lead to an analogous increase in error.

**Figure 3.**
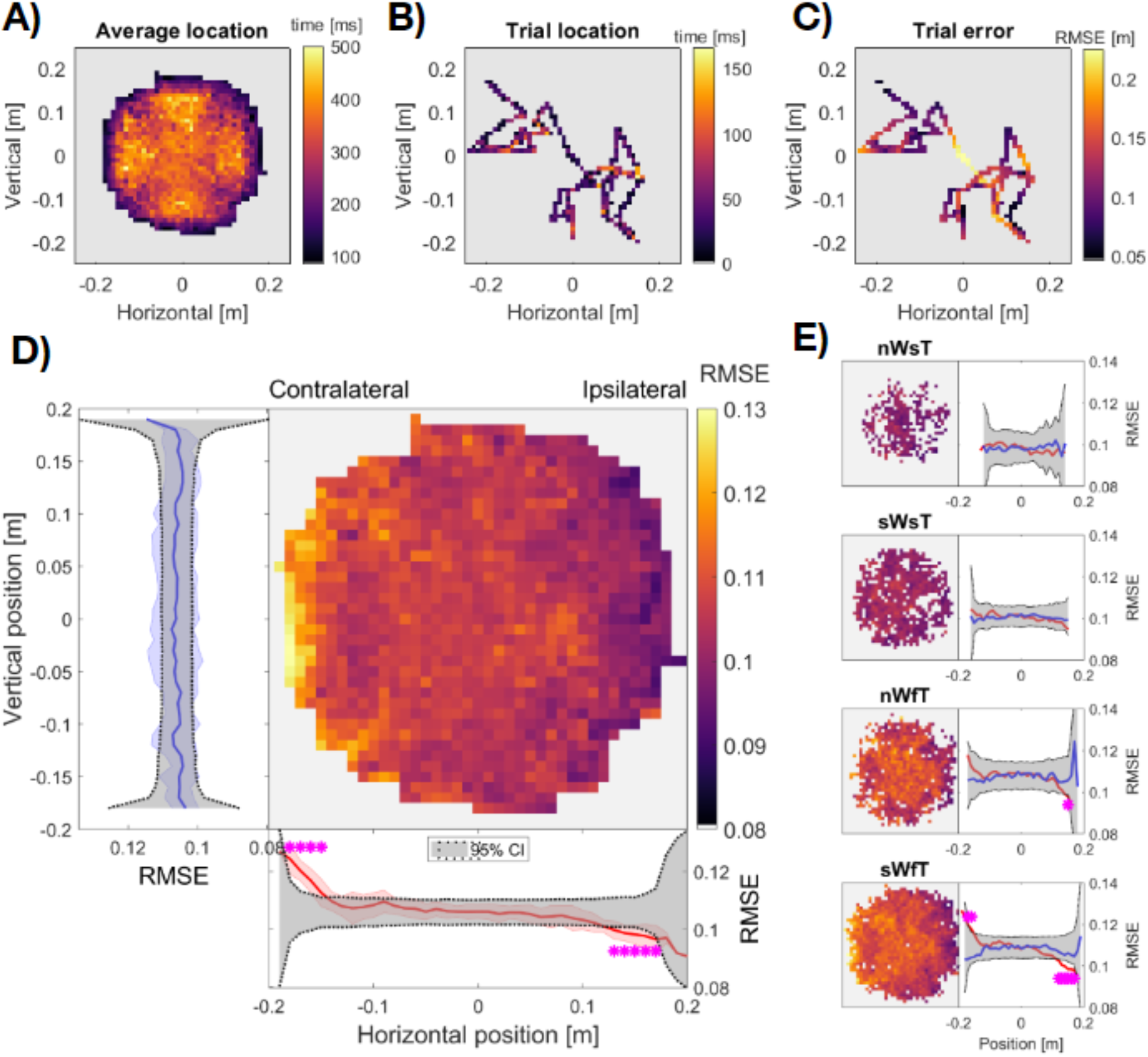
Distribution of target locations and reaching errors within the target motion boundary. **A)** Grand average target distribution across all participants. The heat map displays the average time at each location per trial. **B)** Single trial example of changes in target location. **C)** Single trial error of the same example trial. The heat map displays RMSE for each location. **D)** Grand average reaching error per location. Average vertical error is shown by the blue trace at left (averaging within all rows). Average horizontal error is shown by the red trace at the bottom (averaging within all columns). Blue and Red shading shows the SD across rows and columns, respectively. Dark grey shaded regions in **D)** display the 95% Confidence Interval of error calculated from location-permuted data and magenta asterisks show significant target error compared to this null distribution (p < .05, FDR corrected). **E)** Heat maps show grand average error per condition (n/sW, normal/slow Walk; s/fT, slow/fast Target). Note that the colour scale is truncated and that light grey indicates RMSE < .08, to show the change in relative error across spatial locations. Single traces to the right show average error along one dimension. Vertical is shown in blue, horizontal in red, with 95% CI per trial type in dark grey.

To quantify these changes in error, we calculated the average error on either the horizontal or vertical dimension, and compared the observed error at each horizontal or vertical position with a null-distribution resampled from all locations (**Figure 3D**, see Methods). This analysis revealed that performance was significantly worse when reaching into contralateral peripersonal space (*p* < .001), and significantly improved on the ipsilateral side (*p* < .001), compared to the 95% CI of error sampled from all locations. There were no significant changes in error when reaching along the vertical dimension.

Given the asymmetry in reaching errors was in the horizontal axis, we analysed whether this difficulty reaching accurately into contralateral space was mediated by whether participants were supported by their left or right foot. We hypothesised that if the contralateral error was driven by kinematic or postural demands, then this difference may be larger when swinging the foot ipsilateral to the reaching hand (cf. **Figure 2H**). Contrary to this expectation, there was no difference in error across peripersonal space based on stance (**Supplementary Figure 1**). Upon visual inspection, the effect on the contralateral side was not present in each walking speed x target speed condition and was strongest in the most challenging condition: slow walk with fast target (sWfT: **Figure 3E**). The data in this condition show that walking slowly increases the spatial distribution of reaching error overall and especially when reaching into contralateral peripersonal space. The reduced ability to reach contralaterally arises despite the greater postural stability afforded by a slower more conservative gait, a result we return to in our Discussion.

### The phases of locomotion rhythmically modulate reach error

We have shown that overall reach error is greatest when walking slowly, and when reaching to the contralateral side. We next delved further into this overall error by performing an analysis to quantify whether error changed over the step-cycle. For this, we used Euclidean error (jointly determined by the vertical, horizontal and depth dimension) as a measure of tracking performance and all conditions were examined. We also compared the absolute reach error on each movement dimension, and time-course of reach error when supported by the left or right foot..

The time-course of Euclidean reach error is shown in **Figure 4H**. While the overall pattern of **Figure 2** is preserved (i.e., higher error when walking slowly, tracking fast targets), the difference in reach error when walking at slow vs fast speeds is shown to oscillate, reaching a maximum at the beginning of the swing-phase of each step. We statistically evaluated these error differences with a series of paired-sample *t*-tests at each time-point in the step-cycle. For slow-target conditions, the difference in error when walking at different speeds began immediately prior to footfall, and was protracted through the first half of each step (*clusters* 1st – 45th, 79th – 100th percentiles, *p*_FDR_ < .05). For fast-target conditions, differences in error based on walking speed were confined to a shorter period (*cluster* 14th – 32nd percentile, *p*_FDR_ < .05).

**Figure 4.**
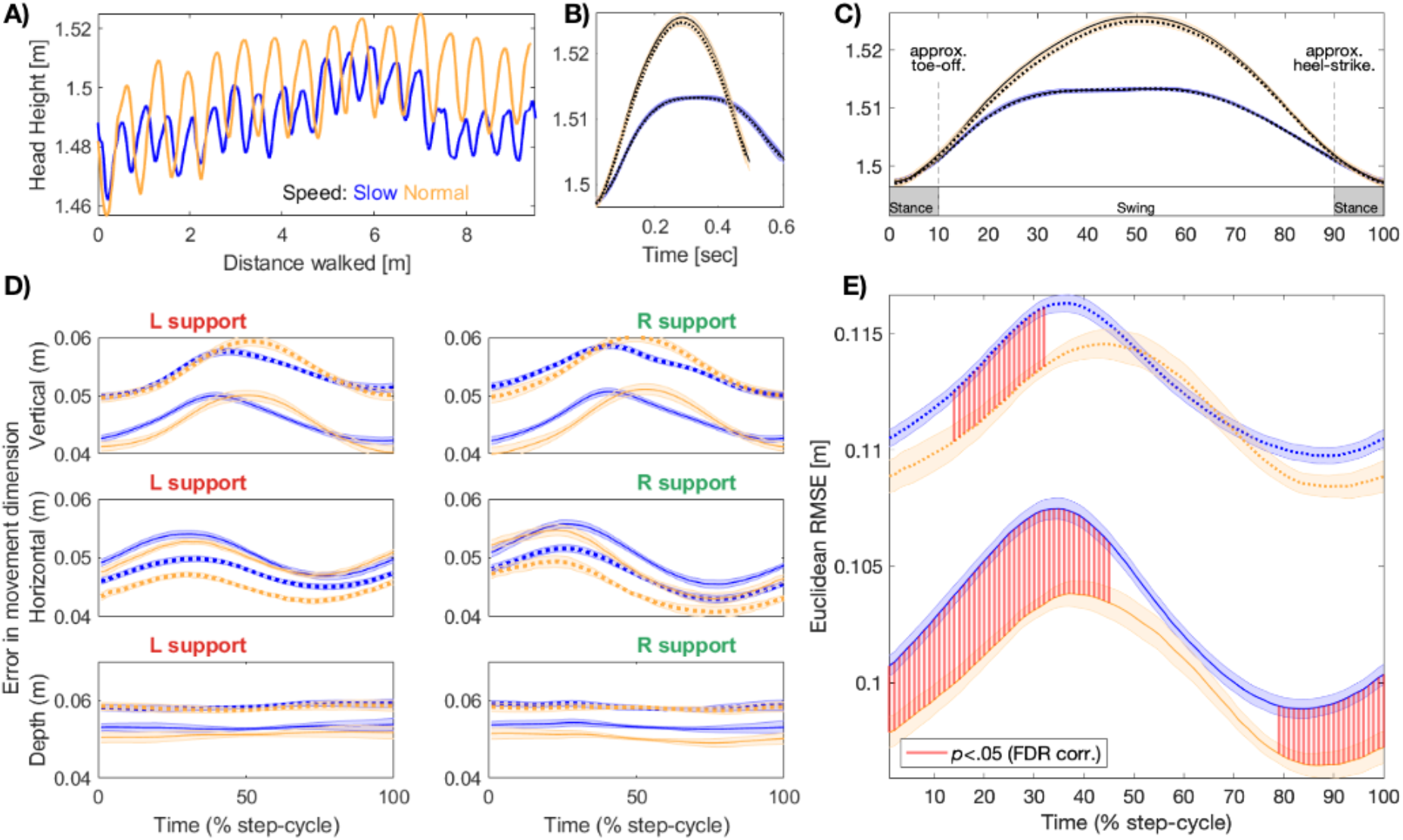
The phase of locomotion modulates reach error. **A)** An example of head height plotted over time for a single trial. Each trial involved walking along a 9.5 m path at either normal (yellow) or slow speed (blue). **B)** Grand average head-height over the step-cycle, epoched per condition (yellow: normal walking speed; blue: slow speed). Solid lines show head-height during slow target conditions, broken lines for fast target conditions. Shading represents mean and ±1 SEM. Note all four conditions are displayed, yet target speeds closely overlap. **C)** Average head-height resampled between 1-100% of step-cycle. Labels showing stance and swing phases of the step-cycle are based on prior research (see Methods). The same resampling procedure was used in **D-E)** to investigate error over the step-cycle. **D)** Average absolute error over the step-cycle on the vertical (top row), horizontal (middle row) and depth axis (bottom row), when supported by the left or right foot (columns) respectively. **E)** Euclidean error over the step-cycle. Red shading represents significant differences between walking speeds, within a target speed condition. Large differences (not visualised) remained between target speed conditions (p < .05, FDR corrected).

As a complement to the Euclidean error, we also quantified fluctuations in error magnitude upon the vertical, horizontal, and depth axes separately, finding distinct patterns in each dimension across the phase of locomotion. Vertical error was modulated roughly symmetrically over the step-cycle, peaking around the midpoint of each step before falling to a baseline level around the time of footfall. In contrast, horizontal error was sinusoidally modulated, and peaked in the ascending phase of the step-cycle, regardless of left- or right-foot support. Error in the depth dimension was relatively constant, presumably due to the smooth and predictable linear motion of our walking guide. **Figure 4D** displays the time-course of reaching error over the step-cycle for each of these conditions.

Together, these results demonstrate that condition-average differences in target tracking based on target walking speed are not stationary, with phasic modulations and optimal periods of sensorimotor precision over the step-cycle.

### Cross-correlogram analysis: Walking slowly increases reaction times

We performed a series of cross-correlogram (CCG) analyses to serve as a proxy for reaction time over the step-cycle. Cross-correlograms estimate the correlation between two time-series over multiple time-points, by shifting the time lag (temporal offset) between them. Following previous research (Bonnen et al., 2017; Cormack et al., 2015; Mulligan et al., 2013), we computed CCGs of the velocity traces of each time-series, and focused on the time lag at which the correlation peaked in each condition (**Figure 5**).

**Figure 5.**
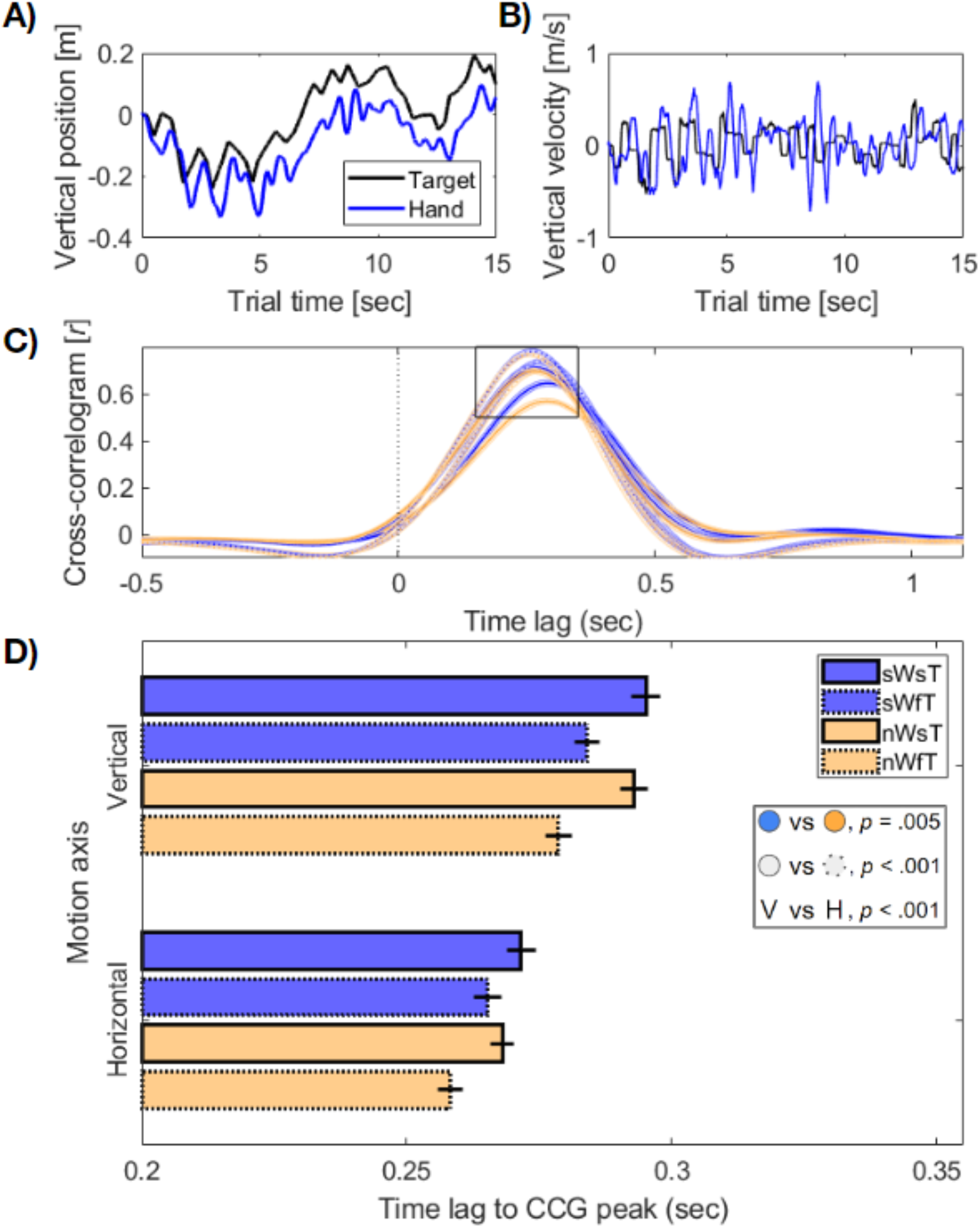
Cross-correlogram (CCG) workflow and results. Examples of single trials from one participant showing **A)** position on the vertical axis, and **B)** the velocity change of the same trials used to compute CCGs. **C)** Shows the overall cross-correlogram, averaged across participants within trial types. A-C show data for the vertical dimension only, identical analyses were performed for the horizontal dimension. **D)** Mean time-lag to the peak in CCG function per trial type. Slow/Normal walk speed denoted by blue/yellow colours, and slow/fast target speed by solid/broken lines. Error bars show ±1 SEM corrected for within-participant comparisons. Legend (inset) summarises significant main effects.

We measured the CCG lag separately on the horizontal and vertical axes, and observed a pattern of reaction times that indicated a speed-accuracy trade-off for fast-targets. Specifically, although distance-based error was larger in these conditions (cf. Figure 2 and 4) reaction times were also faster than when tracking the slower moving targets (**Figure 5**). Participants were also faster overall when walking at normal speeds. We statistically evaluated the difference in peak CCG lag between conditions in a 2×2×2 ANOVA (walk speed x target speed x axis). Significant main effects were observed for walking speed (*F*(1,24) = 9.51, *p* = .005, η_p_^2^ = .29), and target speed (*F*(1,24) = 40.87, *p* < .001, η_p_^2^= .63). We also observed a main effect of motion axis (*F*(1,24) = 49.18, *p* < .001, η_p_^2^ = .67), which indicated that responses were slower overall when correcting for position shifts on the vertical axis. There was additionally an interaction between target speed and motion axis (*F*(1,24) = 8.49, p=.001,η_p_^2^ = .26), such that the difference in reaction times between target speeds was greater for vertical movements. **Figure 5** displays a summary of these results.

### Reaction time varies over the step-cycle

After observing that accuracy fluctuated over the step-cycle (**Figure 4**), we next turned to whether reaction-times also changed based on the relative phase of locomotion. Accordingly, we applied a windowed cross-correlogram analysis (wCCG), to assess whether the lag in peak CCG function varied over the step-cycle. The wCCG method focuses on a short sliding window, and was originally developed to account for dependencies between behavioural time-series which may not be stable over time (Boker et al., 2002; Roume et al., 2018). Here, we calculated wCCG functions between the time-series for hand and target velocities, for both the vertical and horizontal axes, and assigned wCCGs to a relative position in the step using a three step process (see Methods for details). In brief, we first computed the wCCG within a short, sliding window of 111 ms, with 11 ms overlap. As a second step, each wCCG was matched to its simultaneously occurring step, and the wCCG function was allocated to a bin representing relative position in that step-cycle. Each wCCG functions was allocated to a step quintile (i.e., for when the centre of each sliding window fell within the 1-19%, 20-39%, 40-59 % 60-79%, and 80-99% quantiles of a single step, respectively) and averaged within each quintile. The comparison of their time-lag to wCCG peak then served as a proxy for changes in reaction-time over the step-cycle.

Our analyses revealed significant changes in wCCG lag over the step-cycle (**Figure 6**). There was a significant interaction between motion axis and step quintile (*F*(4,96) = 6.32, *p* < .001, η_p_^2^ = .21), indicating that time-lag was affected by position in the step-cycle, but differently on the horizontal and vertical motion axes. As a result, we performed post-hoc analyses within either the vertical or horizontal direction of motion. Our post-hoc analyses revealed that on the vertical axis of motion, reaction times were fastest in the second quintile, during the ascending phase of the step-cycle (*F*(4,96) = 12.53, p < .001, η_p_^2^ = .34; paired-sample *t*-tests, *p*-values correcting for a family of 10; quintile 1 vs quintile 4, *t*(24) = - 3.59, *p* = .009; q2 vs q4, *t*(24) = −6.21, *p* <.001; q2 vs q5, *t*(24) = −4.50, *p* = .001; q3 vs q4, *t*(24) = −4.86, *p* < .001; q3 vs q5, *t*(24) = −4.05, *p* = .004). All other comparisons between quintiles were non-significant (*p*s > .05). Similarly, on the horizontal axis of motion (*F*(4,96) = 3.39, *p* = .012, η_p_^2^ = .12), post-hoc analyses revealed that reaction times were fastest at the beginning of the swing phase of each step, before increasing throughout the step-cycle until the approximate heel strike (q1 vs q4, *t*(24) = −3.46, *p* = .020).

**Figure 6.**
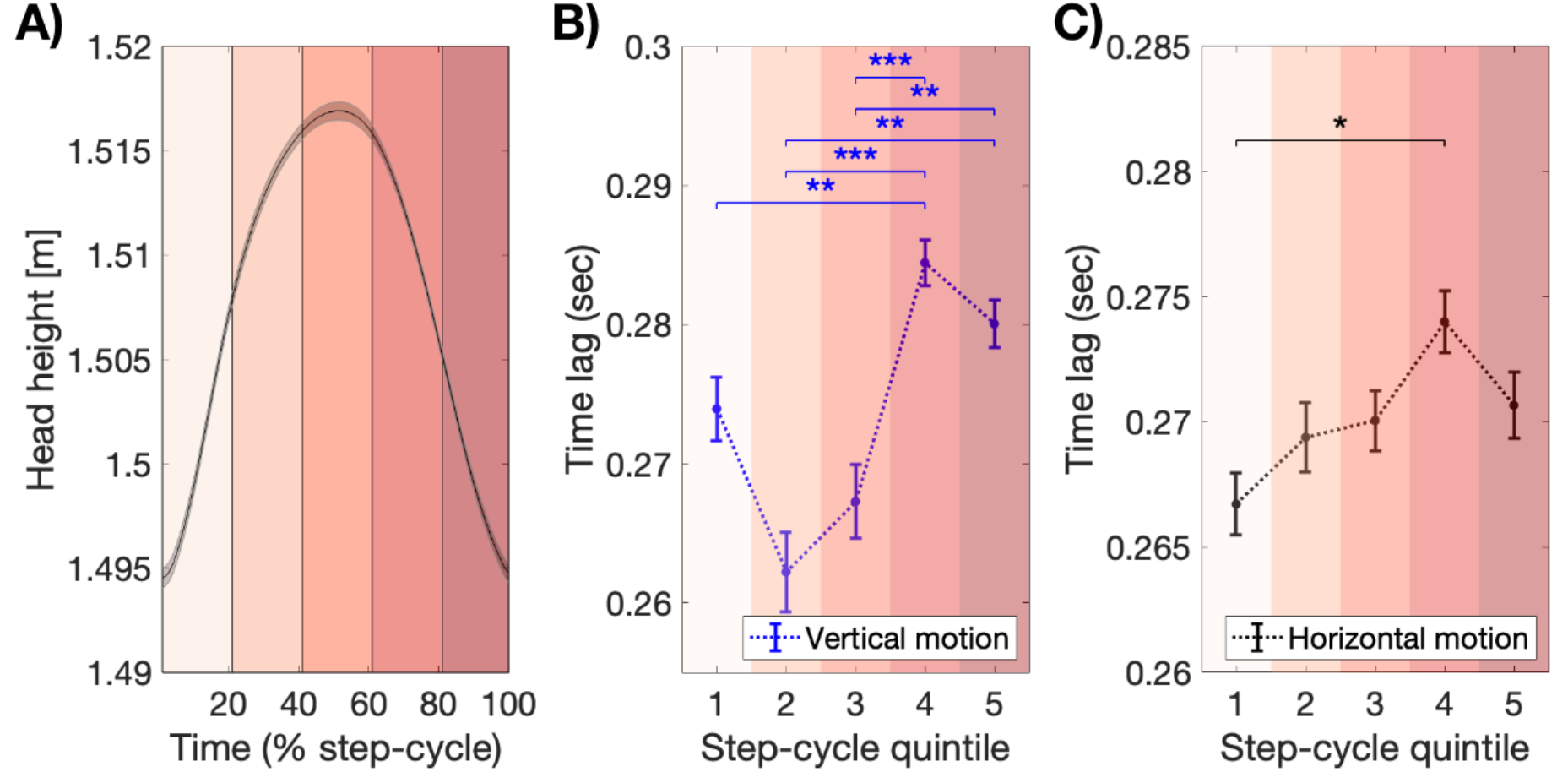
Reaction-time modulates with phase of the step-cycle. **A)** Average head height over all steps. Our wCCG analyses sorted CCG lag into step quintiles shown in shades of Red. **B)** Time lag for the peak in wCCG correlation, averaged per step quintile, shown separately for vertical (blue) and **C)** horizontal (black) motion tracking. In both cases, reaction-times were significantly modulated by the phase of locomotion, fastest in the swing-phase of each step, before slowing prior to footfall. *** p < .001; * p < .05. For wCCG per trial type, see **Supplementary** Figure 2.

In sum, these analyses show that the spatio-temporal tracking speed of a target is fastest during the swing phase of each step, before slowing around the preparation time for the next step, prior to heel strike.

## Discussion

We combined motion-tracking, continuous psychophysics and wireless virtual reality to investigate how steady-state human locomotion affects sensorimotor performance. We operationalised sensorimotor performance by computing accuracy and reaction time measures from a continuous reaching task, and found that normal walking speeds were advantageous. Reach error and reaction times both improved when walking at a normal pace, in contrast to our expectations based on prior research. We also observed phasic modulations to both reach accuracy and reaction time during locomotion. We interpret these results with reference to recent work highlighting that the step-cycle imposes rhythmic changes to sensory processing, potentially due to the ballistic demands of motor-preparation and periods of increased vestibular demand within each step (MacNeilage & Glasauer, 2017).

### Decreased tracking error and faster reaction times when walking naturally

We found that tracking of both fast and slow targets benefitted from walking at a normal pace. Specifically, average tracking error (in Euclidean space), and response times derived from cross-correlograms were both affected by our instruction to walk at a slower speed. These results are in contrast to previous research showing that a more conservative gait is adopted to enable challenging prehension movements (Carnahan et al., 1996; Nashner & Forssberg, 1986; Rinaldi & Moraes, 2015; van der Wel & Rosenbaum, 2007), and that slower walking is often adopted in dual-task scenarios (Al-Yahya et al., 2011). We also observed that reach error slightly increased when standing on the ipsilateral (right) foot, when walking at normal speeds. This effect of foot-support echoes previous results, where either the ipsilateral foot (Cockell et al., 1995; Rinaldi & Moraes, 2015), or contralateral foot (Carnahan et al., 1996; van der Wel & Rosenbaum, 2007) were favoured during reach at walk termination, in a task-dependent manner.

Our task differs from previous designs by combining continuous walking and reaching within peripersonal space. By not enabling participants to adjust their walking speed, we reason that cognitive or biomechanical interactions may be responsible for the counterintuitive finding of increased reach error at slower walking speeds. First, it is possible that our instruction to maintain a slower than normal walking pace imposed unexpected cognitive demands upon primary task performance. Critically, our walking speeds were not self-determined, and prior research has shown that human participants struggle to maintain smooth and slow oscillatory movements that are externally paced (Park et al., 2017). Pedalling at a non-preferred cadence has also been shown to decrease measures of attention (Akaiwa et al., 2022), and dual-task reaction times slow in young adults when asked to walk slowly (Lajoie et al., 1993, 2016). Steady-state locomotion is increasingly being recognised to impose demands on executive function, and attention (Woollacott & Shumway-Cook, 2002; Yogev et al., 2005; Yogev-Seligmann et al., 2008), leaving the possibility that attentional demands increased when walking slowly to an externally imposed speed within our task (Nascimbeni et al., 2015).

Aside from cognitive demands, biomechanical constraints may also have limited reach performance when walking slowly. Previous research has identified a temporal coupling between limb-movements, such that motor commands may be programmed in an integrated or whole-body fashion (Kelso et al., 1979; Marteniuk et al., 2000; Marteniuk & Bertram, 2001; Rosenbaum et al., 2001). Indeed, changes in the difficulty of a prehension task can affect gait, suggesting an interdependence between upper and lower limb movements (van der Wel & Rosenbaum, 2007). Previous work has also found evidence for increased wrist velocity during reach when walking, compared to stationary conditions (Carnahan et al., 1996), where it was suggested that the speed of arm swinging during locomotion may mediate reach behaviour. Here we can extend this comparison to the speed of the arm swinging at different walking speeds. Similar examples of interlimb coupling have been shown between left and right hands when executing simultaneous reaching tasks (Kelso et al., 1979).

Whether limited by biomechanics or cognitive demands, our results suggest that walking slowly is detrimental to reach performance in a continuous tracking task. In addition to these condition based differences, distinct alterations to reach performance also occurred over the step-cycle, at both walking speeds.

### Optimal accuracy and reaction times during distinct phases of locomotion

As a complement to the speed-dependent change in reach accuracy outlined above, we also observed that reach error was dictated by the phase of the step-cycle, during both normal and slow walking conditions. This is in contrast to some previous work, which has described how relative to an end-point in world-space, the trajectory of reaching behaviour is identical whether standing or walking (Marteniuk et al., 2000; Marteniuk & Bertram, 2001). This ‘motor equivalence’ was taken as evidence that reach behaviour was capable of perfectly compensating for the postural changes introduced by walking, as a control strategy to cope with reaching in dynamic contexts. In our data, reach performance was relative to an end-goal in world space (the moving target), yet clear oscillations in error occurred during the step-cycle, in contrast to the motor equivalence hypothesis (Figure 4). More specifically, reach error was largest during the first-half of each step, in the approximate swing-phase, and decreased prior to the time of heel-strike. These phasic changes were present in each walking speed and target speed combination. We also observed that reaction times, computed from the lag-to-peak in windowed cross-correlogram functions, were fastest for movements early in the step-cycle (Figure 6). We note that while small, the maximum difference in reaction times over the step-cycle was 22 ms (*vertical motion,* quintile 2 vs 5), a similar magnitude to the facilitatory effects of spatial cueing on visual detection (Chica et al., 2014).

Our evidence of phasic changes to reach performance over the step-cycle extends prior work investigating the initiation of discrete upper-limb movements during locomotion. Carnahan et al. (1996) recorded the envelope of upper-limb electromyographic (EMG) activity when walking alone, and compared this activity to combined walking with reaching, and standing and reaching conditions. Arm movements were fastest when participants were walking, and reach movements during walking were superimposed upon the profile of EMG activity recorded during walking only conditions. This overlap was taken as evidence that reach behaviour was entrained to the normal rhythm of the step-cycle, to capitalise on the oscillatory movement of upper limbs during normal walking. Earlier work has also found that voluntary and discrete upper limb actions are preferentially initiated at specific phases of the step-cycle. Nashner and Forssberg (1986) observed that treadmill walking participants preferred to initiate voluntary arm-pulls around the time of heel-strike, avoiding upper limb movements when in the swing phase of each step. Similarly, Muzzi et al (1984) observed that the voluntary initiation of clapping was dictated by the step-cycle, showing a strong entrainment to heel-strike times. When reaching for a stationary object, Rinaldi and Moraes (2015) observed that reaching was initiated after ipsilateral heel-contact, and superimposed upon the existing wrist kinematics of locomotion. In combination, the alignment of past research showing preferential initiation of discrete upper limb movements around the time of heel-strike fits well with our measures of reaction-time derived from continuous tracking behaviour. This alignment of upper and lower limb movements may be to capitalise on the conservation of momentum, and to maintain postural stability (Graci, 2011; Matthis & Fajen, 2013; Meyns et al., 2013). Although walking feels relatively smooth and continuous, the ballistic nature of movement preparation appears to mediate the efficiency of upper-limb control. Recent work has also shown that visual information necessary to plan each step must be received in a critical window prior to heel-strike for smooth locomotion to occur - further emphasising the ballistic nature of sensory processing during locomotion (Matthis et al., 2017, 2018).

The timing of performance changes within the step-cycle are noteworthy for a number of reasons. Maximum reach error occurred following toe-off, in the early swing phase of the step-cycle. Reaction times were at their fastest following toe-off, and slowed in the approach to heel-strike of each step. Past research has quantified that head movements are least predictable at these stages of the step-cycle, predicting an increased reliance on vestibular signals during these periods (MacNeilage & Glasauer, 2017). In support of this prediction, prior research has also demonstrated how vestibular signals play a more critical role in maintaining balance and posture during these times (Bent et al., 2004; Dakin et al., 2013). It is plausible that the changes in reach performance we have demonstrated are partly driven by this sensori-motor reweighting. Whether other tasks less reliant on balance are similarly impacted over the step-cycle will be an important area of research to understand the impact of changing vestibular demands on other aspects of perception.

### Increased tracking error in contralateral peripersonal space

Our high-resolution position tracking enabled us to additionally compare how reach performance was executed across peripersonal space. We found that error increased at the left-extrema of the target’s motion boundary, and was lowest on the ipsilateral side. No equivalent change in error was apparent along the vertical axis, despite an equal target distance from the starting point of each trial. Our follow-up analyses confirmed that this increase in contralateral space was not driven by left/right foot support, and increased with a more conservative gait during the slow-walk conditions.

An increase in the distribution of reach error during slow walk conditions is noteworthy, as typically, a more conservative gait is adopted to enable reach-to-grasp behaviour. As outlined above, it is possible that our instructions to participants to maintain a slower than normal pace imposed additional cognitive or mechanical demands which were superimposed upon their task performance. Indeed, the largest spatial asymmetry in reach error was observed in the challenging slow walk, fast target condition (cf. Figure 3E). One possibility is that if externally posed slow-walk conditions did increase cognitive demands, this demand could have produced the rightward shift in attentional bias previously reported in visuo-spatial tasks (Bellgrove et al., 2004; Manly et al., 2005; Pérez et al., 2009). Future research may explore this possibility, by investigating whether the horizontal asymmetries we have reported in reach accuracy are accentuated with further dual-task demands, or dissipate when participants walk at a self-selected slower than normal pace.

### Future research

Future research may also simultaneously measure other co-occurring bodily rhythms over the step-cycle. Eye-movements (Cao et al., 2020; Matthis et al., 2018), heart-rate (Kirby et al., 1989; Niizeki et al., 1993, 1996; Niizeki & Saitoh, 2014), and respiration (Bernasconi & Kohl, 1993; Bramble & Carrier, 1983; H.-T. Lee & Banzett, 1997) all show coupling to walking behaviour - leaving open the possibility that the variations in reach behaviour we report also co-occur with these autonomous bodily functions (Glass, 2001).

An additional possibility is that the timing of eye-movements may contribute to changes in reach error. In our paradigm, we deliberately deployed a small target which would not obscure our participant’s field of vision, yet the possibility remains that eye-movements made to plan future footfall could have coincided with a change in reach error. Similarly, future research may benefit from whole-body position tracking to more accurately establish the precise phases of the step-cycle (such as heel-strike, toe-off, centre-of-mass position, etc), in order to enable how reach behaviour dynamically impacts upon other gait kinematics.

## Conclusion

In summary, continuous peripersonal tracking performance was poorer when walking slowly, contrary to our expectations based on prior research. Phasic changes to performance were also clear over the step-cycle, with both accuracy and reaction-times optimal prior to the heel-strike of each step. The findings demonstrate that the normal stages of human movement restrict sensory performance, highlighting the importance of studying natural and dynamic behaviour instead of relying on traditional static techniques in psychological research.

## Supplementary Figures

**Supplementary Figure 1.**
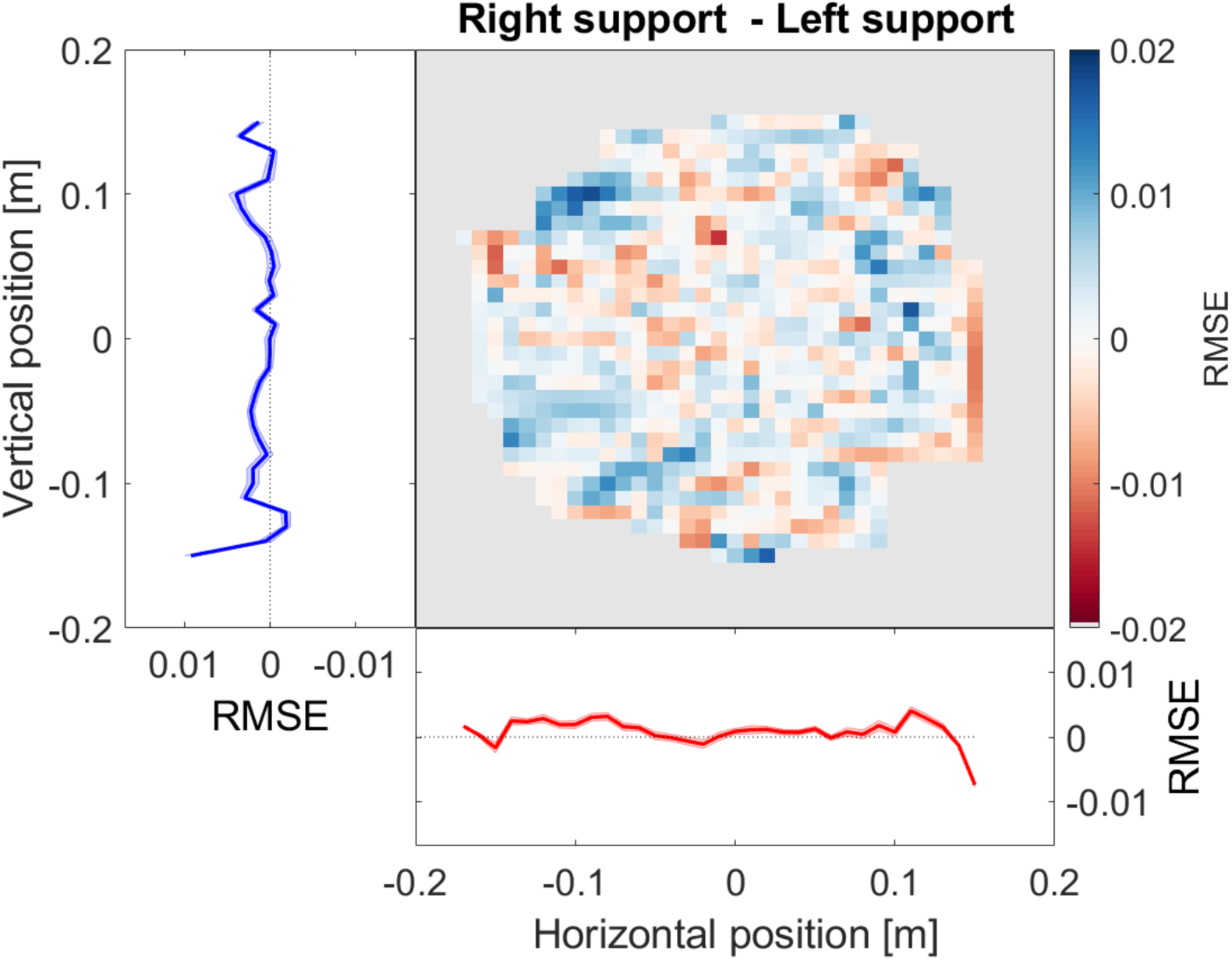
No difference in peripersonal reach error when balanced on the left or right foot. All panels plot the difference in RMSE when comparing right support to left support stance. The heat map displays RMSE for each location. Left) Average difference in error at each position on the vertical axis (when averaging within all rows). Blue lines and shading display the average ± 1 SEM corrected for within-participant comparisons. Bottom) Average difference in error at each position on the horizontal axis (when averaging within all columns). Red lines and shading as in D. The magnitude of error when comparing left/right foot support, is never significantly different from zero after correcting for multiple comparisons.

**Supplementary Figure 2.**
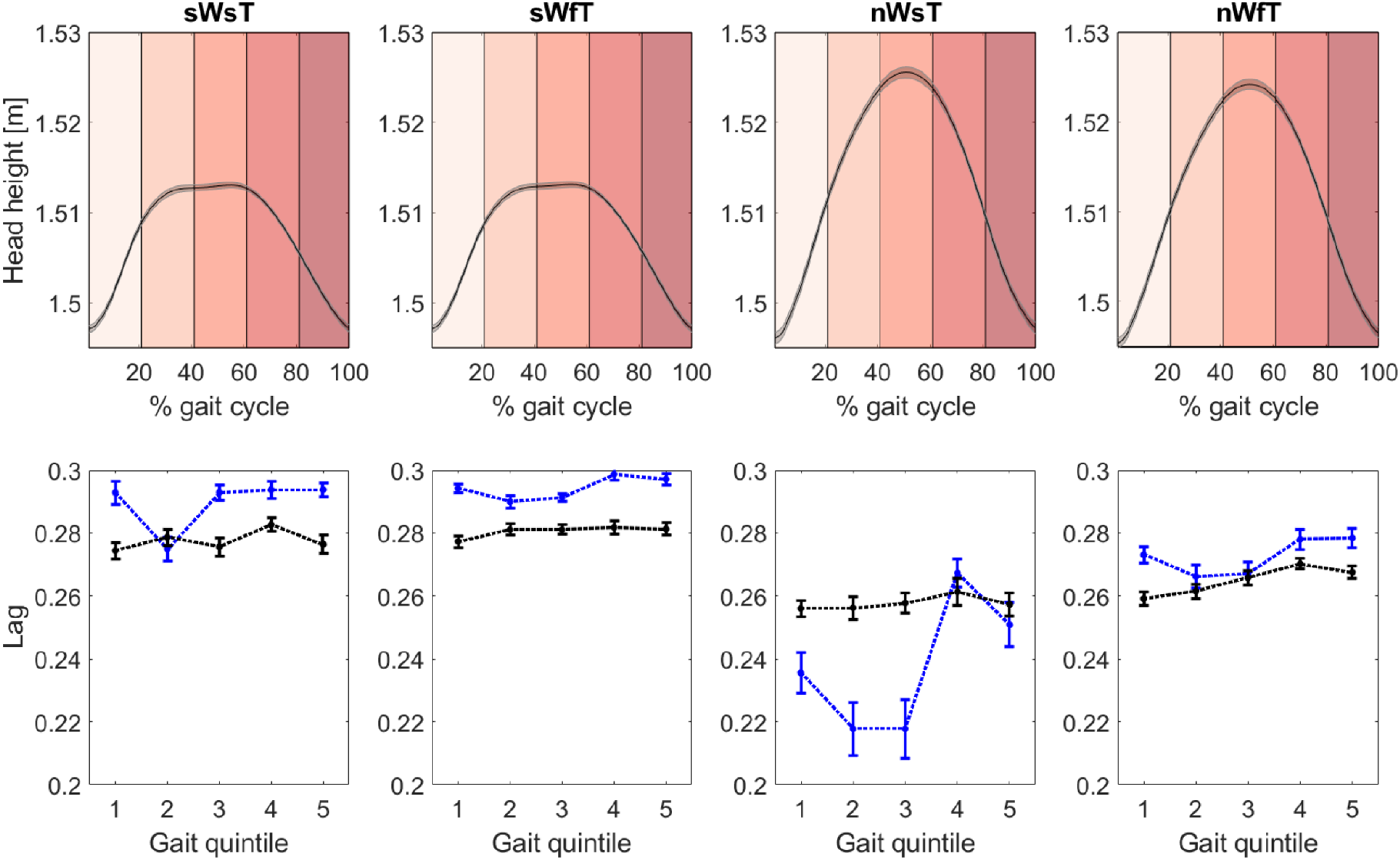
Windowed cross-correlogram per trial type. Blue and Black lines correspond to vertical and horizontal position shifts respectively. Error bars for 1 SEM corrected for within participant comparisons.

